# Sex-biased gene expression in rhesus macaque and human brains

**DOI:** 10.1101/2020.07.17.208785

**Authors:** Alex R. DeCasien, Chet C. Sherwood, James P. Higham

## Abstract

Sexually dimorphic traits (i.e. phenotypic differences between males and females) are largely produced by sex-biased gene expression (i.e. differential expression of genes present in both sexes). These expression differences may be the result of sexual selection, although other factors (e.g., relaxed purifying selection, pleiotropy, dosage compensation) also contribute. Given that humans and other primates exhibit sex differences in cognition and neuroanatomy, this implicates sex differences in brain gene expression. Here, we compare sex-biased gene expression in humans and rhesus macaques across 16 brain regions using published RNA-Seq datasets. Our results demonstrate that most sex-biased genes are differentially expressed between species, and that overlap across species is limited. Human brains are relatively more sexually dimorphic and exhibit more male-than female-biased genes. Across species, gene expression is biased in opposite directions in some regions and in the same direction in others, suggesting that the latter may be more relevant in nonhuman primate models of neurological disorders. Finally, the brains of both species exhibit positive correlations between sex effects across regions, higher tissue specificity among sex-biased genes, enrichment of extracellular matrix among male-biased genes, and regulation of sex-biased genes by sex hormones. Taken together, our results demonstrate some conserved mechanisms underlying sex-biased brain gene expression, while also suggesting that increased neurodevelopmental plasticity and/or strong sexual selection on cognitive abilities may have played a role in shaping sex-biased brain gene expression in the human lineage.

## Introduction

In many species with genetic sex determination, male and female genomes differ by only a few genes located on sex-specific chromosomes. Accordingly, sex-specific phenotypes are largely produced by sex-biased gene expression. Sex differences in gene expression are facilitated by various genomic factors, including relaxed purifying selection, higher tissue specificity (i.e., lower pleiotropy), and imperfect dosage compensation. These differences may also result from sexual selection, a process of evolution by which traits are selected due to their positive effects on mate choice and mating strategies (Darwin 1871). When sexual selection produces sex differences in optimal trait values and the associated underlying gene expression levels, sex-biased expression may emerge as a partial or total resolution to this conflict (Innocenti et al. 2010; Cox and Calsbeek 2009; Ellegren and Parsch 2007). Indeed, both comparative and experimental studies have provided evidence that sexual selection may be linked to observed patterns of sex-biased gene expression. For example, the level of phenotypic sexual dimorphism predicts the proportion of male-biased gene expression in bird gonads and spleens (Harrison et al. 2015), subordinate male wild turkeys exhibit de-masculinized transcriptomes relative to dominant males (Pointer et al. 2013), and enforced monogamy causes gene expression to become feminized in male *Drosophilia* (Hollis et al. 2014). Furthermore, sexual selection pressures can vary greatly between species, and sex-biased genes tend to be differentially expressed between species and to show greater inter-species divergence of both protein-coding sequence and gene expression compared to non-biased genes (Ellegren and Parsch 2007; Grath 2016; Ranz et al. 2003). Although sex-biased gene expression is prevalent in many species, its magnitude varies across tissues and developmental stages (Grath 2016). For example, the gonads exhibit the highest level of sex-biased gene expression across species (Grath 2016; Ellegren and Parsch 2007; Parisi et al. 2004), while the brain varies in its level of dimorphism relative to other tissues (e.g. Yang et al. 2006; Mayne et al. 2016). Furthermore, studies suggest the degree of sex-biased gene expression generally increases during development (Grath 2016; Mank et al. 2010; Magnusson et al. 2011). However, human brains appear to exhibit higher levels of sex-biased gene expression during the prenatal period and puberty than during other developmental periods (Kang et al. 2011; Shi et al. 2016), likely reflecting the organization of sex differences early in development and activation by circulating sex hormones during puberty (Phoenix et al. 1959).

Sexual dimorphism often extends to behavior and cognition, implicating sex differences in neuroanatomy and/or brain gene expression (Mitchell 1980; Jacobs 1996; Knickmeyer et al. 2009; Jazin and Cahill 2010; Ruigrok et al. 2014; Ritchie et al. 2018; c.f. Joel et al. 2015). To date, most studies of sex-biased gene expression in the brain are limited to individual species or regions, or whole brains. Taxa that have been examined include zebrafish (Santos et al. 2008, Wong et al. 2014), chickens (Kaiser and Ellegren 2006), songbirds (two species: Naurin et al. 2011), *Drosophila* (two populations: Catalan et al. 2012), butterflies (five species: Catalan et al. 2018), mice (Rinn et al. 2004; Xu et al. 2002; Dewing et al. 2003; Yang et al. 2006; Li et al. 2007; Armoskus et al. 2014; Naqvi et al. 2019), rats (Naqvi et al. 2019), dogs (Naqvi et al. 2019), marmosets (Reinius et al. 2008), cynomolgus macaques (Reinius et al. 2008; Naqvi et al. 2019), and humans (e.g. Reinius et al. 2008; Vawter et al. 2004; Rinn et al. 2004; Reinius 2009; Kang et al. 2011; Shi et al. 2016; Trabzuni et al. 2013; Werling et al. 2016; Naqvi et al. 2019). Although humans represent the majority of studies on this topic, only one study has explicitly placed human brain sex-biased gene expression in a comparative context. Specifically, Reinius and colleagues (2008) demonstrated that the occipital cortices of humans and cynomolgus macaques, both of which exhibit body size sexual dimorphism indicative of male-male competition, exhibit moderate levels of sex-biased gene expression, whereas expression in the non-size dimorphic marmoset shows almost no sex difference (Reinius et al. 2008). This suggests that sexual selection may have played a role in shaping patterns of sex-biased gene expression in primate brains, similar to studies of other taxa and tissues (e.g. Harrison et al. 2015). Interestingly, humans exhibited even more sex-biased genes than macaques (Reinius et al. 2008), suggesting that brain sex differences may be greater in humans than in macaques even though the latter have higher levels of body and canine size sexual dimorphism. If this is the case, it could reflect stronger sexual selection on cognitive and sensorimotor skills, greater neurodevelopmental plasticity (Sherwood and Gomez-Robles 2017), and/or social experience inequities in humans; however, comparisons of more than one brain region are needed to firmly assess this. In addition, the macaque is the most popular nonhuman primate model for human neurological disorders, many of which show sex differences in prevalence and presentation, so assessing the extent to which sex-biased gene expression is conserved across these species could also have important implications for studies using this model. Here, we address these gaps in our knowledge by examining patterns of sex-biased gene expression in the brains of humans and rhesus macaques across 16 brain regions using published RNA-seq datasets (Zhu et al. 2018). Notably, these publicly available datasets have relatively large sample sizes and a high number of brain structures.

We made a number of predictions grouped thematically (Table 1).

**TABLE 1.**

| TABLE 1 |  |  |  |
| --- | --- | --- | --- |
| PREDICTIONS | JUSTIFICATION | METHOD(S) | SUPPORTED? |
| **General patterns** |  |  |  |
| [1] Samples will cluster strongly by species, region, and age, and more subtly by sex | While a majority of human brain structural MRI scans can be correctly classified according to sex using various discriminant and classification approaches (Del Giudice et al. 2016; Rosenblatt 2016), there is a large amount of overlap between male and female distributions (Joel et al. 2015). Conversely, studies have shown large species, region, and age differences in brain gene expression (e.g. Khaitovich et al. 2004; Kang et al. 2011; Zhu et al. 2019; Naqvi et al. 2019). | Dimensionality reduction (UMAP), Hierarchical clustering | Yes |
| **Species differences** |  |  |  |
| [2] Many sex-biased genes will be differentially expressed between human and rhesus macaque brains | Across species, sex-biased gene expression across a range of tissues tends to be species-specific and differentially expressed between species (Ranz et al. 2003; Ellegren & Parsch 2007). For example, most sex-biased genes are only biased in the brains of one of two songbird species (Naurin et al. 2011), in one of three primate species (Reinius et al. 2008), in one of five butterfly species (Catalan et al. 2018), or in a subset of five mammalian species (Naqvi et al. 2019). | Sex-biased gene identification (DESeq2), Differentially expressed gene identification (DESeq2) | Yes |
| [3] Human brains will exhibit more sex-biased genes than rhesus macaque brains across regions | Sexual selection may have been particularly strong throughout human brain evolution since the emergence of language would have allowed more direct sexual selection on cognitive abilities. Correspondingly, humans exhibit numerous sex differences in neurobiology and the prevalence of certain neurological disorders (e.g. Ruigrok et al. 2014), and there is some evidence that human brains exhibit relatively greater sex-biased gene expression than other taxa. For example, while the mouse brain exhibits less sexually dimorphic gene expression than other tissues (Yang et al. 2006; Li et al. 2017), the human anterior cingulate cortex is one of the most dimorphic non-gonadal tissues in terms of gene expression (Mayne et al. 2016; Gershoni & Pietrovski 2017). In addition, human occipital cortices exhibit more sex-biased genes than those of cynomolgus macaques or marmosets (Reinius et al. 2008). | Sex-biased gene identification (DESeq2) | Yes |
| [4] Across species, shared patterns of sex-biased gene expression profiles will be limited to certain regions and genes | Female rhesus macaques outperform males in spatial memory tasks when landmarks are removed (Herman & Wallen 2007) and have relatively larger corpus callosum volumes (Knickermeier et al. 2009). These sex differences are in the opposite pattern to those found in humans (e.g. DeLacost-Utamsing & Holloway 1992; Sandstrom et al. 1998). | Sex-biased gene identification (DESeq2), Correlations between sex effects | Yes |
| **Species commonalities** |  |  |  |
| **Distribution** |  |  |  |
| [5] Gene expression will be sex-biased in the same direction across most regions within species, with functionally connected regions exhibiting the most similar sex effects | Models of the mouse functional connectome suggest that both anatomical connectivity and correlated gene expression support functional connectivity (Krienen et al. 2016; Mills et al. 2018). Furthermore, studies suggest that when genes are sex-biased in multiple tissues, the bias is likely to be in the same direction across tissues (Naqvi et al. 2019). | Correlations between sex effects | Partially (negative/no correlation for some macaque regions) |
| [6] Cortical areas will be the most dimorphic regions | The majority of studies of sex-biased gene expression in human brain have shown that cortical areas are the most dimorphic brain areas (e.g. frontal and temporal cortex: Kang et al 2011; anterior cingulate cortex: Mayne et al 2016; occipital cortex: Trabzuni et al. 2013). | Sex-biased gene identification (DESeq2) | Partially (also non-cortical for macaques) |
| [7] Sex-biased genes will be less pleiotropic within the brain than non-biased genes | Work on other species and tissues suggest that sex-biased genes are relatively less pleiotropic, allowing them to better respond to narrow selection regimes (Mank et al. 2008; Naqvi et al. 2019). | Tissue specificity analyses | Yes |
| [8] Most brain regions will exhibit more male- than female-biased genes | Sex-biased genes in the brain tend to be male-biased (humans: Reinius et al. 2008; Kang et al. 2011; Shi et al. 2016; cynomolgus macaques: Reinius et al. 2008; mice: Armoskus et al. 2014; birds: Kaiser & Ellegren 2006; Naurin et al. 2011; Drosophila: Catalan et al. 2012) due to sexual selection-driven masculinization of the genome (Harrison et al. 2015; Singh & Kulathinal 2005; Singh & Artieri 2010). | Proportion comparisons (binomial tests) | Partially (humans only) |
| **Function** |  |  |  |
| [9] Male-biased gene expression profiles will be enriched for extracellular matrix organization and energy production processes; female-biased gene expression profiles will be enriched for immune system processes | Previous work suggests that male-biased brain genes are enriched for energy production and extracellular matrix organization (zebrafish: Wong et al. 2014; cynomolgus macaques: Reinius et al. 2008; humans: Reinius et al. 2008; Mayne et al 2016; Shi et al. 2016; Ziats & Rennert 2013), while female-biased genes tend to be enriched for immune-related pathways (humans: Trabzuni et al. 2013; Mayne et al 2016). | Gene ontology enrichment analyses (topGO) | Partially (male-biased extracellular structure only) |
| **Regulation** |  |  |  |
| [10] Sex-biased gene expression will be regulated by sex hormones (e.g. testosterone, estrogen, progesterone) | Androgen and estrogen responsive elements play a role in the regulation of sex-biased brain gene expression (zebrafish: Santos et al. 2008; humans: Trabzuni et al. 2013; Mayne et al. 2016). | Motif enrichment analyses (HOMER) | Partially (macaque enrichment NS) |

## Results

### Samples cluster strongly by species, age, and region (Prediction 1)

Dimensionality reduction analyses suggest that species cluster together within age groups (i.e. fetal versus non-fetal samples; Figure 1). Given that age clearly accounts for a significant amount of gene expression variation, we controlled for age throughout our differential expression analyses and reran analyses on fetal and non-fetal subsets separately (see Methods). Species differences are also apparent, with species showing more differences (i.e. less overlap) at the fetal stage (Zhu et al. 2018; Figure 1). Non-fetal cerebellum and thalamus samples cluster together across species, separating from all other brain regions (Figure S1). By contrast, samples do not appear to cluster by sex (Figure 1). These patterns provide support for Prediction 1, and suggest that sex effects on brain gene expression are more subtle than species, region, or age effects.

**Figure 1.**
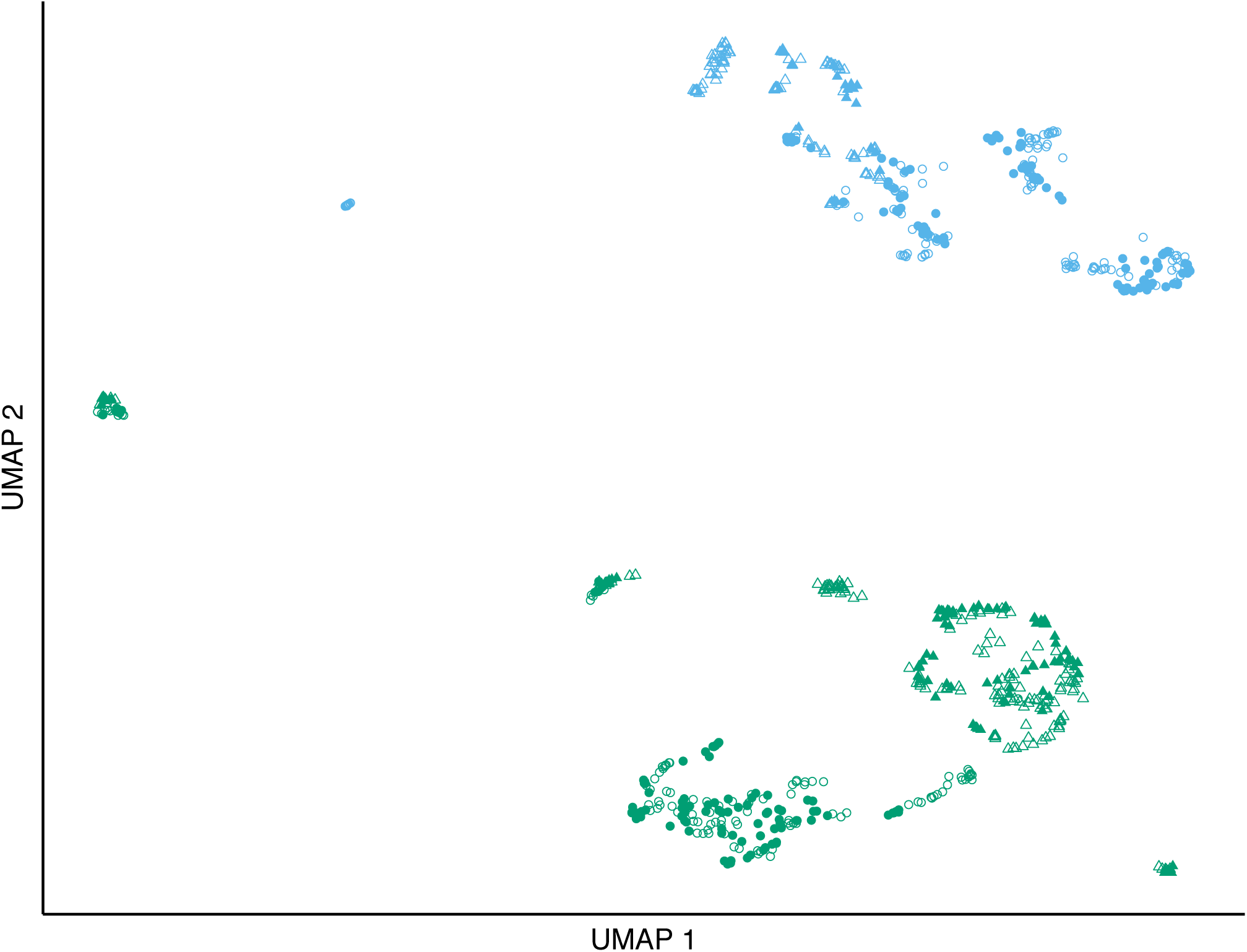
Samples cluster clearly by age and species. Uniform Manifold Approximation and Projection (UMAP) plot of brain gene expression. Each dot represents one sample. Dot color indicates age (blue = fetal; green = non-fetal), shape indicates species (circle = human; triangle = macaque), and fill indicates sex (open = male; sex = female). A UMAP plot depicting regional differences in provided in Figure S1.

Hierarchical clustering analyses suggest that macaque and human samples within the same age group (i.e. fetal or non-fetal) are more similar to each other than to same species samples within a different age group; Figure S2). Within fetal samples, cerebellum samples cluster together as the outgroup to all other regions, consistent with previous reports (e.g. Lo and Hilbush 2001). Within age groups and species, cortical areas tend to cluster together. Samples cluster subtly by sex only in some cases (e.g. some subsets of non-fetal human cortical samples are generally separated by sex: see node 733), in support of Prediction 1. Lack of clustering by sex may reflect very low resolution at mid-levels of the dendrogram (i.e. where AU=BP=0).

### Many sex-biased genes are differentially expressed between species (Prediction 2)

For each species and brain region, we identified hundreds of sex-biased genes using various differential expression (DE) p-value cutoffs (see Methods; Figure 2; Table S1). Fetal samples exhibit more sex-biased genes than non-fetal samples within each species (Table S1). Within both species, the majority of genes exhibiting sex-bias in at least one brain region were also differentially expressed between species in at least one brain region (72-78% within each species across all DE p-value cutoffs; Table S2). This is generally consistent across sample size permutations (mean = 60-61% for both species; Table S3). An even higher proportion was observed in the fetal subset (95-97% within each species across all DE p-value cutoffs; Table S2). Within brain regions, many genes that exhibited sex-bias were also differentially expressed between species within the same brain region, although the proportion varied by region and DE p-value cutoff (humans: 9-43%; macaques: 0-40%; Figure S3; Table S2). This is generally consistent across sample size permutations (range across regions: humans: 0-55%; macaques: 0-53%; Table S4). An even higher proportion was observed in the fetal subset (humans: 35-65%; macaques: 36-73%; Table S2). Overall, our Prediction 2 is supported.

**Figure 2.**
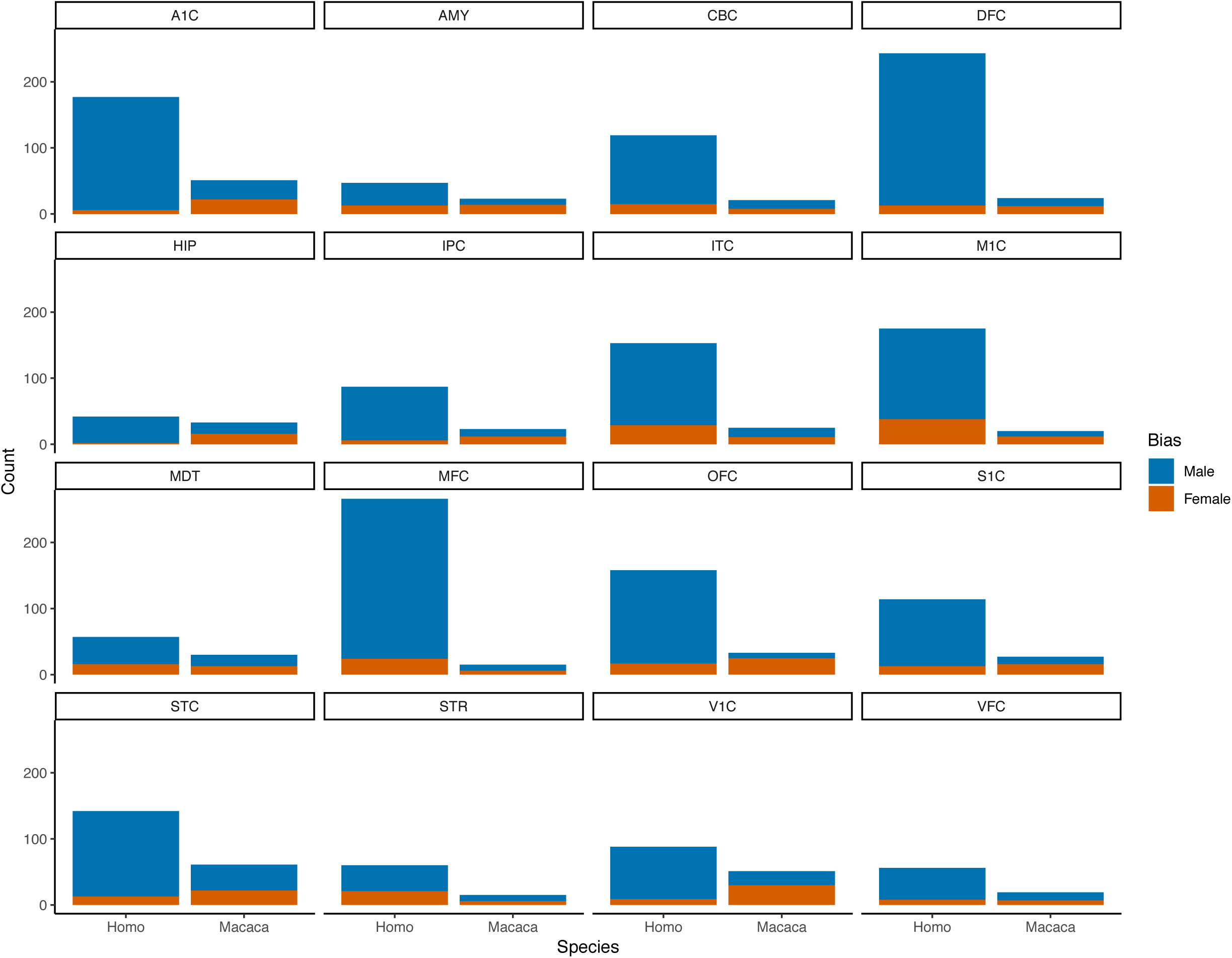
Human brains exhibit relatively more sex-biased genes and display more male-than female biased genes. Bars represent counts of sex-biased genes identified using the p-value cutoff p<0.005 across regions and species. Colors indicate direction of bias (blue = male-biased; orange = female-biased). Counts of sex-biased genes using different p-value cutoffs are available in Table S1.

### Human brains exhibit more sex-biased genes than rhesus macaque brains across regions (Prediction 3)

In support of our Prediction 3, humans exhibit more sex-biased genes than rhesus macaques across the majority of regions and DE p-value cutoffs (Figure 2; Table S1). This difference is particularly pronounced in the MFC and M1C across all DE p-value cutoffs. These results are generally consistent within age groups (Table S1) and across permuted sample sizes (humans exhibit more sex-biased genes than rhesus macaques in 71% of all iterations across all regions).

### Across species, shared patterns of sex-biased gene expression profiles are limited to certain regions and genes (Prediction 4)

Between species, correlations between the sex effects across genes differ by region. There is a significant positive relationship between sex effects (i.e. genes tend to be biased in the same direction across both species) for the STC, VFC, MFC, M1C, S1C, IPC, and CBC (Figure 3; Table S5). There is no relationship between sex effects across species for the DFC, MDT, or STR (Figure 3; Table S5). There is a negative relationship between sex effects (i.e. genes tend to be biased in the opposite direction across humans and rhesus macaques) in the A1C, AMY, HIP, ITC, OFC, and V1C (Figure 3; Table S5). These results are generally consistent across permuted sample sizes (i.e. the mean correlation across iterations is the same sign as in the original analysis), except for the A1C, OFC, and V1C; Table S5). Results vary within age groups, and there are more regions that exhibit a negative correlation for sex effects across species within the fetal compared to the non-fetal subset (Table S5). Notably, interspecific correlations are relatively high for the CBC and IPC across all analyses (Table S5). Overall, these results support our Prediction 4.

**Figure 3.**
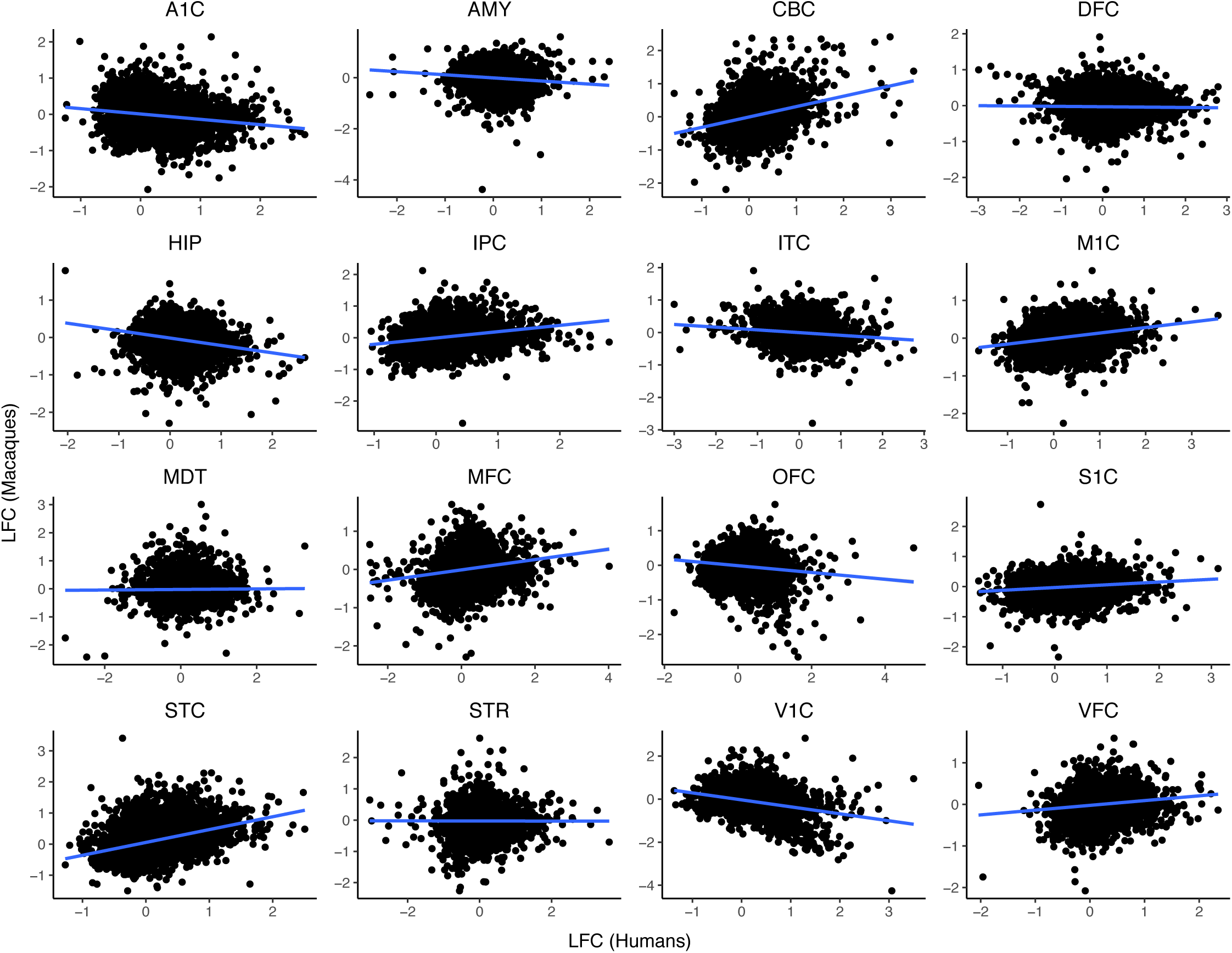
Across species, genes tend to be sex-biased in the same direction in some regions and in the opposite direction in others. Log2 fold change (LFC) values represent the difference in expression between male and female samples. Across genes within regions, LFC values for macaques are regressed on LFC values for humans. Regression lines are in blue. Regression details are provided in Table S5.

The male-biased Y chromosome genes and female-biased *XIST* gene exhibit conserved sex-biased expression across species and regions (Tables S6-S8). Aside from these genes, relatively few genes exhibit conserved, region-specific patterns of sex-bias genes across species (Tables S6-S8). For example, at the most stringent DE p-value cutoff (p<0.005), such genes include: female-biased expression of *PNPLA4* in the DFC, male-biased expression of *ELN* in the STC, male-biased expression of *POSTN* in the A1C, male-biased expression of *ABI3BP* in the S1C, male-biased expression of *NAGA* in the MDT, female-biased expression of *PDYN* in the MDT, and female-biased expression of *KDM5C* in the DFC, VFC, and OFC. Of these genes, only *NAGA, PDYN, KDM5C* were not top-ranked (i.e. prior rank < 0.5) in a broad study of differential gene expression predictability (Crow et al. 2019). Additionally, at this DE p-value cutoff, only 3 genes exhibit opposite patterns of sex-biased expression (Table S6).

Among the non-fetal subset, very few genes exhibit conserved or opposite patterns of sex-bias (Tables S6-S8). Compared to both non-fetal samples and the overall sample, fetal samples exhibit many more genes with opposite patterns of sex-bias and also more genes with conserved patterns (Tables S6-S8). Notable examples that are not likely generic DE genes (i.e. prior rank < 0.5; Crow et al. 2019) include *TFB2M* (male-biased in MFC), *MBIP* (female-biased in STR), *CACNA1I* (male-biased in S1C), *CDH20* (male-biased in MDT), and *PRDX6* (male-biased in MDT), all of which are involved in brain development (Park et al. 2018; Invernizzi et al. 2018; Xu et al. 2018; Matsunaga et al. 2013; Shim et al 2012).

At a less stringent DE p-value cutoff (p<0.05), elastin (*ELN*) and numerous collagen genes tend to be male-biased throughout the human brain; however, these genes tend to be male-biased in only a few regions of the macaque brain (for examples, see Figure S4). This finding is generally consistent within age groups; however, these genes are also likely to be generic DE genes (i.e. prior rank > 0.5; Crow et al. 2019).

### Gene expression is sex-biased in the same direction across most regions within species (Prediction 5)

Inter-regional correlations are generally positive across regions within both species (Figure 4; Table S9). For humans, the sex effect across all genes exhibits a positive correlation between all pairs of brain regions (i.e. genes tend to exhibit the same direction of sex-bias across all brain regions), except for the MFC and STR. For rhesus macaques, the sex effect is positively correlated across the majority of brain region pairs. Negative correlations characterize the following pairs of regions: 1) HIP and STC; 2) CBC and V1C; 3) CBC and OFC. These results are generally consistent across permuted sample sizes (i.e. mean number of region pairs with negative correlations: humans = 5, macaques = 2). Inter-regional correlations tend to be lower among the fetal subsets for both species, with more pairs of regions exhibiting negative correlations (humans = 20; macaques = 5). Overall, these results support Prediction 5.

**Figure 4.**
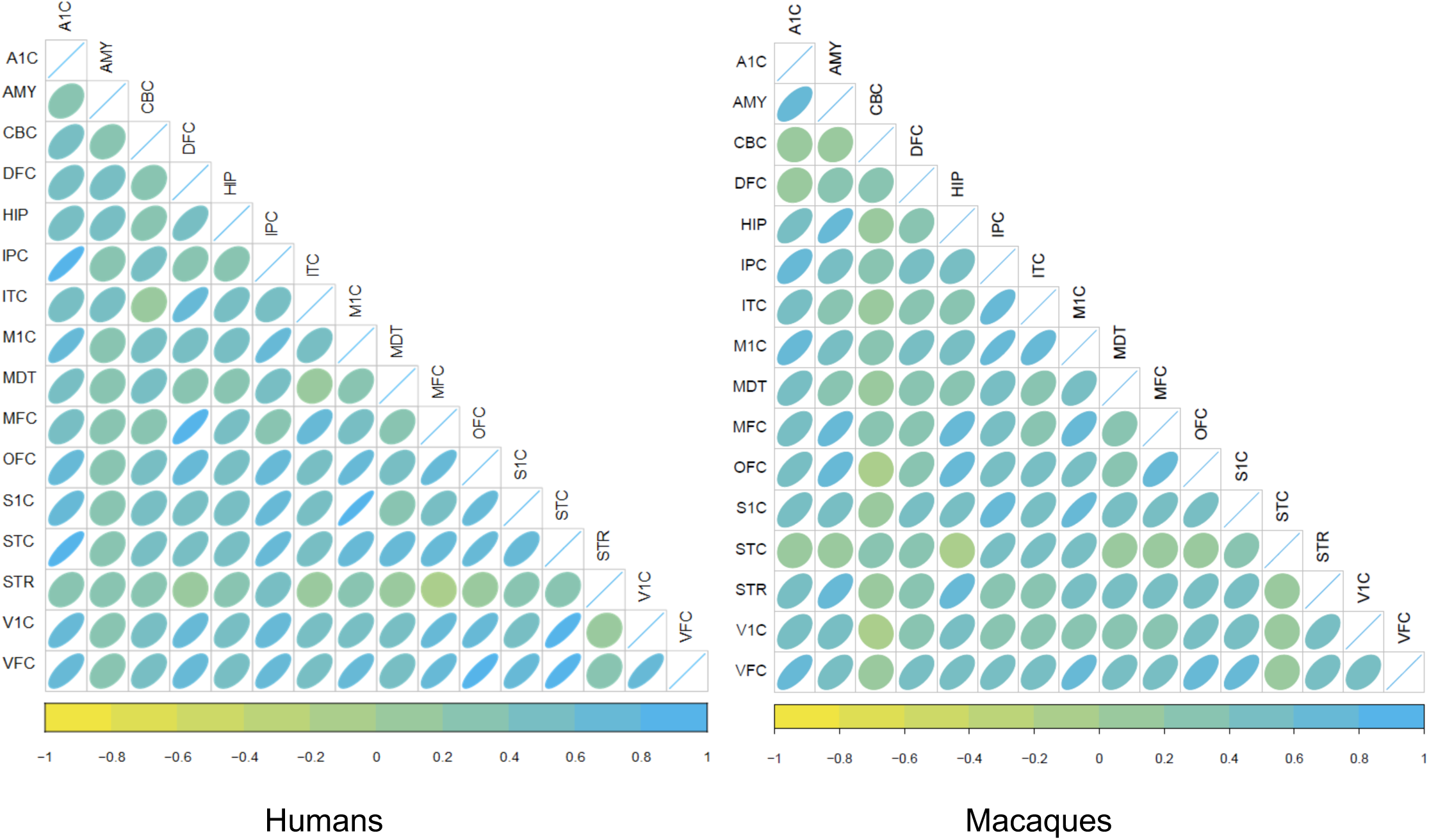
The majority of inter-regional sex effect correlations are positive across both species. Correlation plots of sex effects between all pairs of regions across both humans (left panel) and macaques (right panel). Colors and shapes indicate direction and strength of correlation (more yellow = stronger negative correlation; more green = weak correlation; more blue = stronger positive correlation; circle = weak correlation; oval = stronger correlation in the given direction).

### Cortical areas are generally the most dimorphic regions (Prediction 6)

In support of Prediction 6, human cortical areas (except for VFC) and the cerebellum exhibit more sex-biased genes than subcortical areas (i.e. STR, MDT, AMY, HIP exhibit the lowest number of sex-biased genes) across all DE p-value cutoffs (Figure 2; Table S1). This is slightly different from what is observed for rhesus macaques, as the most dimorphic areas include not only the STC V1C A1C, but also the HIP (Figure 2; Table S1). Interestingly, although the MFC is among the most dimorphic regions in humans, it is among the least dimorphic in rhesus macaques. These results (i.e. cortical areas constituting the most dimorphic regions in humans, but not in macaques) are generally consistent within the fetal and non-fetal age groups (Table S1) and also across permuted sample sizes (Table S1).

### Sex-biased genes are less pleiotropic within the brain than non-biased genes (Prediction 7)

Sex-biased genes exhibit a significantly higher (t-test: p<0.05) mean tissue specificity than non-biased genes across all DE p-value cutoffs. Across species and DE p-value cutoffs, non-biased genes exhibit a mean tissue specificity of ∼0.1, while sex-biased genes exhibit a mean tissue specificity somewhere between ∼0.2-0.3 (see Figure 5). These results are consistent within age groups (i.e., tissue specificity in both fetal and non-fetal subsets is significantly higher among sex-biased genes [mean = ∼0.2] than non-biased genes [mean = ∼0.1]) and also across permuted sample sizes (i.e., sex-biased genes exhibit significantly higher mean tissue specificity than non-biased genes in 98% of human iterations and 100% of rhesus macaque iterations; Table S10). This supports our Prediction 7.

**Figure 5.**
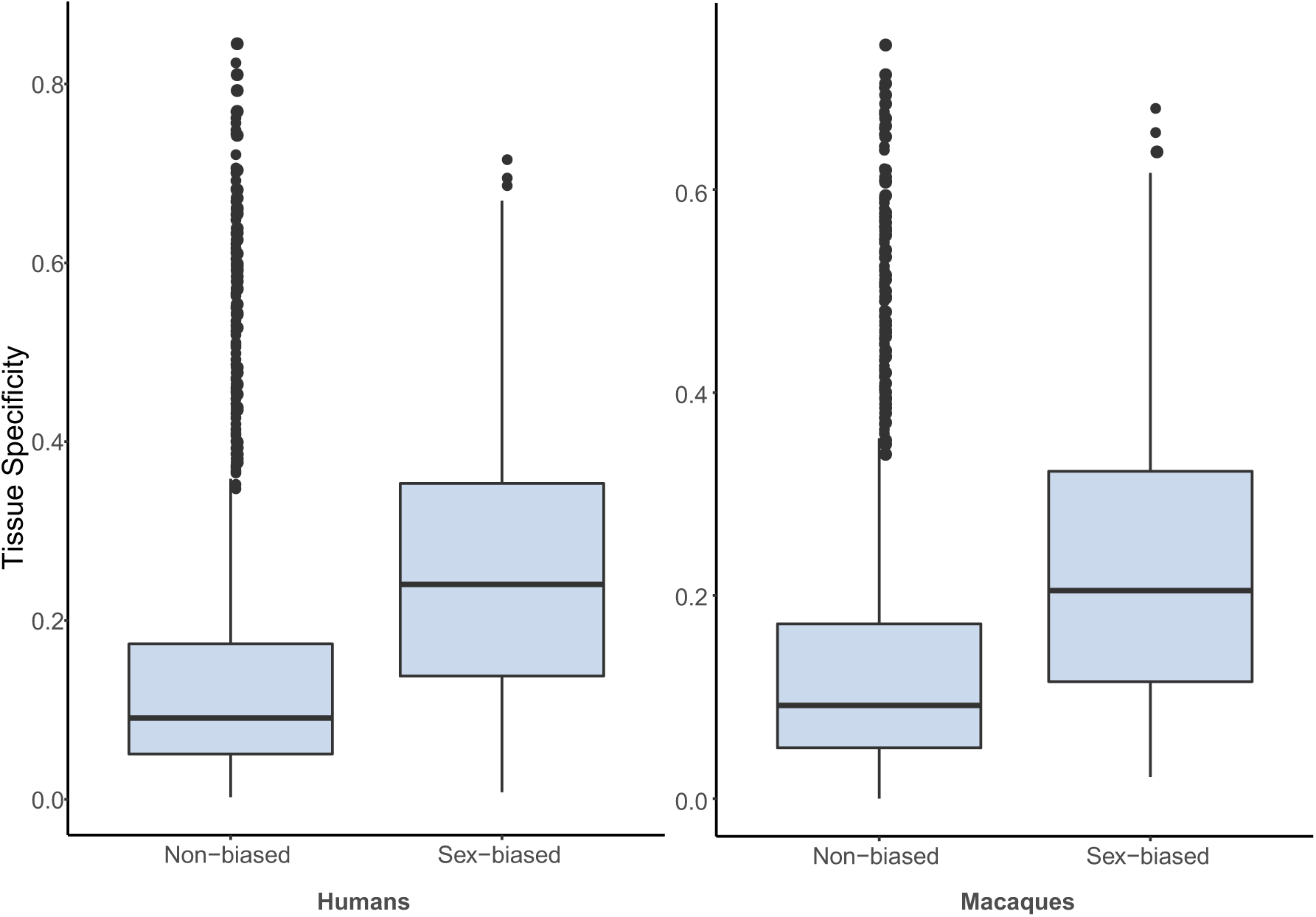
Sex-biased genes exhibit higher tissue specificity than non-biased genes in both species. Box plots show tissues specificity for sex-biased genes (i.e. genes showing bias in any region) and non-biased genes (i.e. all other expressed genes) for both species (horizontal line = median, boxes = interquartile range [IQR], whiskers = 1.5*IQR, dots = outlying points).

### Human but not rhesus macaque brains exhibit more male-biased genes (Prediction 8)

Nearly all human brain regions exhibit significantly more (binomial test: p<0.05) male-biased than female-biased genes at all DE p-value cutoffs (Figure 2; Table S1). This is generally consistent within age groups and across sample size permutations, as only 0-2 regions exhibit significantly more female-biased genes in some analyses (Table S1). The majority of rhesus macaque brain regions do not exhibit a significant difference in the number of male-biased versus female-biased genes, although more regions tend to exhibit significantly more female-biased genes at higher DE p-value cutoffs. This is consistent with analyses of non-fetal macaque samples; however, various regions exhibit significantly more male-or female-biased genes within the fetal subset (Table S1). Overall, these results suggest that Prediction 8 is supported only for human brains.

### Male-biased gene expression profiles are enriched for extracellular matrix organization processes (Prediction 9)

According to the top 30 gene ontology terms derived from each data set, male-biased genes in the brains of both species are enriched (Fisher’s exact test: p<0.05) for extracellular structure organization (Table S11). This is consistent for fetal samples of both species and non-fetal human samples (Table S11). Only male-biased genes in humans are enriched for immune system processes (Table S11). Accordingly, Prediction 9 is partially supported.

### Sex-biased gene expression is regulated by sex hormones (Prediction 10)

The promoter regions of sex-biased genes, compared to those of non-biased genes, tend to be enriched for sex hormone binding sites (Table S12). At the strictest DE p-value cutoff (p<0.005), sex-biased gene promoters in the human brain are significantly enriched for estrogen (p=0.01) and progesterone (p=0.01) responsive elements. Sex-biased gene promoters in the macaque brain exhibit relatively more estrogen responsive elements than background genes, although enrichments are not significant (p=0.10). Fetal samples of both species are significantly enriched (p=0.01) for estrogen, androgen, and progesterone responsive elements (Table S12). Therefore, our Prediction 10 is partially supported.

## Discussion

Overall, this work highlights both distinct and shared features of sex-biased gene expression in the brains of humans and rhesus macaques. Although sex impacts brain gene expression less than age, species, or region, we identified hundreds of genes with sex-biased expression patterns. Given that previous work has shown that sexual selection pressures can shape patterns of sex-biased gene expression, our results suggest that species-specific sexual selection on cognition and sensorimotor skills may have influenced the patterns of sex-biased brain gene expression observed here. We found that sex-biased genes tend to be differentially expressed between species. Additionally, human brains are relatively more sexually dimorphic and exhibit more male-than female-biased genes, with gene expression biased in opposite directions in some regions compared to rhesus macaques. In other brain regions, genes tend to be biased in the same direction across species, suggesting these areas may be better suited for studies of nonhuman models of neurological disorders. The brains of both species share some features relevant to the distribution, function, and regulation of sex-biased genes, including positive correlations between sex effects across regions, reduced pleiotropy among sex-biased genes, enrichment of extracellular matrix structure among male-biased genes, and regulation of sex-biased genes by sex hormones.

Decoupling male and female gene expression can allow sexual selection to act on male and female phenotypes separately. Accordingly, the presence of very different sexual selection pressures across species may partially explain why sex-biased genes tend to be species-specific and exhibit both sequence and expression divergence between species (e.g. Naurin et al. 2011; Harrison et al. 2015; Reinius et al. 2008; Ellegren and Parsch 2007; Grath 2016; Ranz et al. 2003). In fact, a recent study incorporating data from tissue samples from multiple organs across five species suggested that the majority of sex-biased gene expression is not shared between most mammalian species (i.e., sex-biased gene expression tends to be specific to single species or subsets of species), implying these expression patterns arose relatively recently in each evolutionary lineage (Naqvi et al 2019). Another recent study on the brains of five butterfly species had similar results (Catalan et al. 2018). Our results show a similar pattern for the species examined here, since: 1) within age groups, samples generally cluster by age, region and species, rather than by sex; and 2) most sex-biased genes are differentially expressed between species. This pattern is particularly pronounced at the fetal stage in both species. In particular, a greater number of genes exhibit sex bias during the fetal compared to non-fetal stage, and a higher percentage of those genes are also differentially expressed between species. This is similar to previous findings in humans (Kang et al. 2011) and is likely to reflect species differences in the organization of sex differences early in development.

One possibility that is consistent with our data is that sexual selection may have been particularly strong throughout human evolution, leading human brains to be relatively more sexually dimorphic than those of rhesus macaques. While the extent of human neuroanatomical sex differences is debated (e.g., Joel et al. 2015; Del Giudice et al. 2016), there is a general consensus regarding humans sex differences in the prevalence of certain neurological disorders (e.g., autism: Werling et al. 2013), which may be driven by differences in neuroanatomy (Baron-Cohen et al. 2005; Joel 2011) and brain gene expression (Werling et al. 2016). Notably, the emergence of language may have allowed more direct sexual selection on mental faculties, which otherwise could have only been assessed by directly observing relevant behaviors. Brain dimorphism may also be facilitated by evolutionary increases in neurodevelopmental plasticity (Sherwood and Gomez-Robles 2017). This suggests that human brain development is sensitive to the surrounding environment, so neurobiological sex differences in humans are likely to be mediated by systems of inequity in how girls and boys are treated throughout development (i.e., the strong sexual dimorphism in human brains might be a feature of societal exposure rather than because of evolutionary selection). Our results support these ideas, since there are more sex-biased genes in the human brain than in the rhesus macaque brain across all regions. The observed differences between species in non-fetal samples may be further heightened by the laboratory environment of the rhesus macaques, whose opportunities to participate in sex-specific reproductive strategies or other behaviors are limited. In addition, other studies have identified neuroanatomical sex differences in humans that are not observed in the rhesus macaque, such as relatively larger amygdala volumes in males (Knickermeyer et al. 2009). We also found that most sex-biased genes in the human brain are male-biased, while there tends to be no significant difference in the proportion of male- or female-biased genes in the rhesus macaque brain. Across bird species, masculinization (i.e. the proportion of male-biased genes) of gonad and spleen genomes is positively correlated with multiple measures of sexual selection, including relative testis weight, sperm number, and sexual ornamentation (Harrison et al. 2015). Accordingly, these results may reflect relatively strong sexual selection on human cognition, and implicate major evolutionary changes to transcription factors, cis-regulatory sites, and/or post-transcriptional regulators.

In line with species differences in the relative proportion of male- and female-biased genes, genes tend to be biased in opposite directions in some regions across species. Specifically, we found a negative relationship between sex effects in the rhesus macaque versus human A1C, ITC, V1C, OFC, HIP, and AMY. This pattern is particularly pronounced during the fetal stage, during which more regions exhibit negative correlations. This is likely explained by opposite patterns of sexually dimorphic cognitive and sensory abilities, resulting from species-specific sexual selection pressures. For example, as described in Table 1, rhesus macaques and humans exhibit opposite patterns of sex differences in spatial memory tasks (Herman and Wallen 2007; Sandstrom et al. 1998) and corpus callosum volumes (DeLacost-Utamsing and Holloway 1982; Knickermeyer et al. 2009). Additional species differences in expression involve collagen and elastin genes, which tend to be male-biased throughout the human brain but generally only male-biased in a few regions of the rhesus macaque brain. Given that these genes tend to be differentially expressed across human studies (Crow et al. 2019), it is difficult to associate them with a particular phenotype; however, the species differences highlighted here are suggestive of biological significance. Collagens are involved in brain development and may play a role in the development of Alzheimer’s and other dementias (Cheng et al. 2009), which disproportionately affect women (Mazure and Swendsen 2016). Notably, Kang et al. (2011) previously reported sex differences in expression and exon usage for these genes in multiple areas of the human brain. Although these species-specific patterns of sex-biased gene expression suggest that nonhuman models may not effectively represent human sex differences (Naqvi et al. 2019), relevance may vary across tissues and brain regions. For instance, we found that in some regions, including the STC, VFC, MFC, M1C, S1C, IPC, and CBC, gene expression tends to be biased in the same direction across species. Accordingly, these areas may be better suited for model studies of human neurological disorders.

Although numerous differences exist between these species in the patterns of sex-biased brain gene expression, they do share multiple characteristics relevant to the distribution and function of sex-biased genes, in addition to underlying regulatory mechanisms. Within both species, we found that there tends to be a positive correlation for sex effects across regions, in line with previous findings across tissues (e.g. Naqvi et al. 2019). There are more pairs of regions that exhibit negative correlations during the fetal stage. This is likely to reflect higher inter-regional transcriptional differences during this period (Kang et al. 2011). Given that both anatomical connectivity and correlated gene expression support functional connectivity (Mills et al. 2018), we predicted that functionally linked regions would exhibit the highest interregional correlations. In fact, we found that among the most similar regions for both species are the M1C and S1C, areas that modulate each other’s activity to fine tune somatosensation and movements (Borich et al. 2015). In humans, sex effects in the STC are highly correlated with those in the A1C and the VFC, areas which are functionally connected. Initial detection of auditory information occurs in the A1C, with detailed analysis occurring in the posterior superior temporal gyrus. There is a pathway connecting posterior temporal gyrus and vlPFC (Garell et al. 2012), the latter of which processes the level of novelty and allocates attentional resources accordingly (Schonwiesner et al. 2007). In rhesus macaques, the highest interregional correlation is between the HIP and AMY, between which reciprocal connections exist (Saunders et al. 1988) to facilitate processes such as fear conditioning (Phillips and LeDoux 1992).

In both species, cortical areas are the most dimorphic, although the HIP is among the most dimorphic regions in rhesus macaques. In humans, the most dimorphic areas are higher-order cognitive areas (prefrontal areas DFC and MFC) and the primary motor cortex (M1C). Interestingly, the precentral gyrus (containing the M1C) exhibits the most female biased relative volume in humans, and contains a relatively high density of estrogen and androgen receptors (Goldstein et al. 2001). In rhesus macaques, the most dimorphic areas are involved in social cognition (STC), sensory information processing (V1C, A1C), and spatial cognition (HIP). This may reflect “neocorticalization” of cognitive function in humans, resulting from expansion of the human neocortex and, consequently, its enhanced connectivity and more dominant role in overall brain function (Deacon 1990; Streidter 2005). Broadly, these patterns suggest sexual selection has mostly occurred on first-order cognition (i.e. sensory perception) in rhesus macaques and higher-order cognition in humans. We also found that sex-biased genes are less pleiotropic across the brain than non-biased genes. This confirms previous work on non-neural tissues (e.g. Mank et al. 2008) and is the first study to demonstrate this pattern across areas of the same organ. Heightened tissue specificity among sex-biased genes reflects the fact that more narrowly expressed genes tend to evolve relatively fast (Meisel 2011; Mank et al. 2008) since genes with fewer, more specific functions can more easily respond to selection. Interestingly, genes expressed in the brain may exhibit higher tissue specificity than genes expressed in other non-sexual tissues (Li et al. 2017; Ma et al. 2018), which may facilitate sex-biased gene expression in the brain.

In addition to similarities in regional distribution, sex-biased genes in the brains of both humans and rhesus macaques exhibit some similarities in function and regulation. Specifically, we found that male-biased genes are enriched for extracellular matrix structure in both species. Enrichment is not driven by Y chromosome genes; rather, it is due to male-biased expression of multiple autosomal genes, including *POSTN, ELN, ABI3BP*, and collagen genes, across multiple brain areas. Collagens and elastin represent the majority of macromolecules in the extracellular matrix (Frantz et al. 2010), while POSTN secretes an extracellular matrix protein involved in axon regeneration and neuroprotection (Matsunaga et al. 2015), and ABI3BP promotes cellular senescence and extracellular matrix interactions (Delfin et al. 2019). Given that extracellular matrix plays a major role in the development and maintenance of synapses, our results suggest selection for similar sex differences in synaptogenesis and synaptic plasticity across species (Ferrer-Ferrer and Dityatev 2018). In fact, human males have a higher synaptic density than females throughout the cortical layers of the temporal neocortex (Alonso-Nanclares et al. 2008). We should note, however, the extracellular matrix gene module tends to be generically differentially expressed across human studies (Crow et al. 2019). Finally, we found that the promoters of sex-biased genes tend to be enriched for estrogen and progesterone responsive elements. Previous studies of the human brain found significant enrichment of androgen, but not estrogen, responsive genes; however, this result may have been influenced by the fact that the majority of women included were post-menopausal (Trabzuni et al. 2013). Across human tissues, about one third of autosomal sex-biased genes contain androgen or estrogen hormone response elements, suggesting that the majority of sex-biased genes are not under direct influence of sex hormones (Mayne et al. 2016). This is not surprising given that numerous pathways have been identified through which sex hormones indirectly regulate gene expression (e.g. Dyson and Wright 2016).

Taken together, our results demonstrate the potential for both lability and consistency of sex-biased gene expression in the brain across primate species. Previous work on other tissues has indicated that sex-biased genes can exhibit rapid evolutionary turnover (Ranz et al. 2003; Zhang et al. 2007; Harrison et al. 2015). Our work represents an example of such turnover, as human and rhesus macaque brains exhibit major differences in both the number and direction of sex-biased genes, with gene expression in some regions exhibiting overall opposite patterns of sex bias across species. These differences may reflect strong sexual selection on human cognitive abilities, evolutionary increases in neurodevelopmental plasticity, and/or sex differences in societal experiences. Some brain regions show similar sex-specific patterns of expression, implicating conserved or convergent sexual selection mechanisms on associated cognitive functions. Similarities across species in the distribution, function, and regulation of sex-biased genes provide additional evidence that, sex-biased genes are under less evolutionary constraint (Mank et al. 2008), male-biased genes exhibit similar functions across species, and sex hormones play a major role in the regulation of sex-biased genes.

## Materials and Methods

### Data collection

Gene expression data were downloaded from http://evolution.psychencode.org/ (Zhu et al. 2018). Please refer to the original publication for complete details on dissection, RNA extraction, and sequencing methods. To summarize briefly, transcriptomes were generated for tissue samples representing hippocampus (HIP), amygdala (AMY), striatum (STR), mediodorsal nucleus of thalamus (MDT), cerebellar cortex (CBC), and 11 areas of the neocortex (primary auditory, A1C; dorsolateral prefrontal, DFC; inferior posterior parietal, IPC; inferior temporal, ITC; primary motor, M1C; orbital prefrontal, OFC; primary somatosensory, S1C; superior temporal, STC; primary visual, V1C; ventrolateral prefrontal, VFC; and medial prefrontal, MFC) from 36 humans (21M/15F; aged 56 days – 40 years; fetal: 10M/7F; non-fetal: 11M/8F) and 26 macaques (18M/8F; aged 60 days – 11 years; fetal: 7M/2F; non-fetal: 11M/6F). Metadata for individuals included in this study are provided in Table S13 and summarized in Figure S5. Reads were mapped to the human (*Homo sapiens*, hg38) and rhesus macaque genomes (*Macaca mulatta*, rheMac8), respectively, using STAR (Dobin et al. 2013). The XSAnno pipeline was used (Zhu et al. 2014) to generate annotation of human-macaque orthologs, based on human Gencode v25. The XSAnno pipeline incorporated whole genome alignment and local alignment with multiple filters. This resulted in a common annotation set of 27,932 orthologous mRNAs.

To make human data comparable across ages, we: 1) excluded transient structures, including the medial ganglionic eminence (MGE), lateral ganglionic eminence (LGE) and caudal ganglionic eminence (CGE); 2) excluded samples spanning the fetal motor and parietal somatosensory cerebral wall (MSC), parietal cortex (PC) and temporal cortex (TC); and 3) equated the occipital part of the cerebral wall (OCC) and the primary visual cortex (V1C), the dorsal part of the thalamic anlage (DTH) and the medio-dorsal thalamus (MDT), and upper (rostral) rhombic lip and the cerebellum (CBC) (see Li et al. 2017 for details).

### Statistical analyses

#### Dimensionality reduction (Prediction 1)

The Uniform Manifold Approximation and Projection (UMAP) method (McInnes et al. 2018), performed by the umap function in R, was used for dimensionality reduction. UMAP is a non-linear dimensionality reduction method that preserves more of the global structure than other non-linear methods (i.e. t-SNE), and is therefore more appropriate for gene expression data than other methods (e.g. principal components analysis (PCA) is limited to linear dimensionality reduction). Prior to dimensionality reduction, gene counts were converted to counts per million (CPM) and genes with mean CPM less than 2 were removed.

#### Hierarchical clustering (Prediction 1)

Hierarchical cluster analyses were performed by the pvclust function in R (Suzuki and Shimodaira 2006). Correlation was used as the distance measure. This function provides both approximately unbiased (AU) *p*-value and bootstrap probability (BP) value. AU values are calculated using multiscale bootstrap resampling, while BP values are calculated by the ordinary bootstrap resampling (Suzuki and Shimodaira 2006). Prior to hierarchical clustering, gene counts were converted to CPM and genes with mean CPM less than 2 were removed.

#### Identification of sex effects and sex-biased genes (Predictions 2-10)

Differentially expressed genes were computed with the R package DESeq2 (Love et al. 2014). Count data were modeled with a negative binomial distribution to address overdispersion. Size factors were calculated that take into account the total number of reads in different samples, and a dispersion parameter was determined for each gene to account for biological variation between samples.

Intra-species sex-biased genes were computed for each brain region. Within each region, only genes with mean CPM greater than 2 were included in differential gene expression analyses. Generalized linear models of the negative binomial family were fitted for each gene. The fitted models were as follows:

- Rhesus macaques: ∼ sex + age + extraction method
- Humans: ∼ sex + age

Individual variability and moderate sample size are likely to affect the power to observe statistical significance. Given that we wished to include all genes with a propensity for sex-biased expression, we followed similar studies (e.g., Werling et al. 2016) and identified genes with both fold difference magnitudes > 1.2 and unadjusted p-values < 0.05, 0.01, or 0.005 (i.e. DE p-value “cutoffs”) as differentially expressed by sex.

#### Inter-species differentially expressed gene identification (Prediction 2)

Differentially expressed genes were computed with the R package DESeq2 (Love et al. 2014) as above. Inter-species differentially expressed genes were computed for each brain region. Within each region, only genes with mean CPM greater than 2 were included in differential gene expression analyses. Generalized linear models of the negative binomial family were fitted for each gene. The fitted model was:

- ∼ species + age + extraction method

Genes with adjusted p-values < 0.1 were identified as differentially expressed by species.

To test whether sex-biased genes also tend to be differentially expressed between species, we compared: 1) lists of genes identified as sex-biased in any region with genes identified as differentially expressed any region; and 2) lists of genes identified as sex-biased with genes identified as differentially expressed within the same regions.

#### Interspecies comparisons of sex-biased genes (Prediction 3)

The total number of sex-biased genes (male-biased + female-biased) was compared within regions across species at each DE p-value cutoff.

#### Correlations between sex effects within regions across species (Prediction 4)

We examined correlations between the sex effects per gene within each region between species. This was done using linear models (lm function in R). We excluded Y chromosome genes and *XIST* since extreme sex differences in the expression of these genes skew the models towards a positive correlation. Slope estimates were examined to determine the direction of the relationship. Regression models with adjusted p<0.05 (using the Benjamini & Hochberg (1995) correction) were considered significant.

#### Correlations between sex effects across regions within species (Prediction 5)

We examined correlations between the sex effects per gene across all pairs of regions within each species. This was done using correlation analyses (cor function in R). We excluded Y chromosome genes and *XIST* since extreme sex differences in the expression of these genes skew the models towards a positive correlation. Slope estimates were examined to determine the direction of the relationship.

#### Regional comparisons of sex-biased genes (Prediction 6)

Within species, the total number of sex-biased genes (male-biased + female-biased) was compared across regions at each DE p-value cutoff.

#### Tissue specificity analyses (Prediction 7)

Following previous studies, we calculated tissue specificity per gene by calculating τ (Yanai et al. 2004; Mank et al. 2007). Studies suggest this is the most robust method for detecting tissue specificity (Kryuchkova-Mostacci and Robinson-Rechavi 2017). Data for each gene was standardized to CPM. In the formula below, N is the number of tissues examined and CPMmax is the highest expression level detected for a given gene over all tissues examined. The value of τ ranges from 0 to 1, with lower values indicating an expression pattern that is evenly distributed through all tissues examined and higher τ values indicating more variation in expressional levels across tissues and a greater degree of tissue specificity. We conducted a t-test of tissue specificity values between sex-biased and non-biased genes to examine differences in mean tissue specificity. Significant differences in tissue specificity were identified when p<0.05.

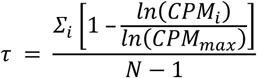

#### Proportion comparisons (Prediction 8)

To test for significant differences in the number of male-versus female-biased genes within each data set (per species and region), we used two-sided binomial tests (binom.test function in the R package stats). Adjusted p<0.05 (using the Benjamini & Hochberg (1995) correction) were considered significant.

#### Gene ontology enrichment analyses (Prediction 9)

Gene ontology (GO) enrichment analyses were performed using the topGO R package (Alexa and Rahnenfuhrer 2010) for sex-biased genes identified using the DE p-value cutoff p<0.005. We used Fisher’s exact tests, which are based on gene counts. Enrichments with unadjusted p< 0.05 were considered significant, as suggested by the package’s creators (Alexa and Rahnenfuhrer 2010). For each data set, we extracted the top 30 terms according to score. The parent child algorithm (Grossman et al. 2007) was used since it determines overrepresentation of terms in the context of annotations to the term’s parents. Other approaches to measuring overrepresentation of GO terms cannot cope with the dependencies resulting from the structure of GO because they analyze each term in isolation. The parent child approach reduces the dependencies between the individual term’s measurements, and thereby avoids producing false-positive results owing to the inheritance problem.

#### Motif enrichment analyses (Prediction 10)

We used HOMER (Hypergeometric Optimization of Motif EnRichment) to analyze the promoters of genes and look for motifs that are enriched in our target gene promoters relative to other promoters. For each species, three target sets were created, which included genes that were either male-biased or female-biased in at least one region across the 3 DE p-value cutoffs. Three background sets were created, which included all non-biased genes per DE p-value cutoff (i.e. genes expressed in at least one region that were not included in either the female- or male-biased gene sets). We searched for motifs from −2000 to +2000 relative to the TSS. Since we only focused on six motifs related to sex hormones (see Table S12), unadjusted enrichment p-values less than 0.05 were considered significant.

#### Subgroup analyses

Given that dimensionality reduction and hierarchical clustering analyses suggest strong separation between fetal and non-fetal samples (see Results), all of the analyses described above (except for dimensionality reduction and hierarchical clustering) were repeated within each age group (i.e. fetal and non-fetal) within each species. Due to time and processing constraints, motif enrichment analyses were only repeated using the most stringent DE p-value cutoff for identifying sex-biased genes (i.e. p<0.005).

#### Sample size effects

Possible effects of unbalanced sample sizes across species and sexes were evaluated using permutations. Specifically, the species/sex group with the minimum number of individuals was the macaque female group, which included samples 2 fetal and 6 non-fetal individuals. Given that samples from all regions are not available for all individuals, this created some sample size variation across regions at each iteration, as in the original, complete data set. At each of 100 iterations, 2 fetal and 6 non-fetal individuals were randomly selected for each species/sex group and the following analyses were repeated using this subset: identification of sex effects and sex-biased genes (using the most stringent DE p-value cutoff, p=0.005), inter-species differentially expressed gene identification, interspecies comparisons of sex-biased genes, correlations between sex effects within regions across species, correlations between sex effects across regions within species, regional comparisons of sex-biased genes, tissue specificity analyses, and proportion comparisons.

## Acknowledgements

We thank Dr. Nenad Sestan, Dr. Andre Sousa, and Dr. Ying Zhu for feedback on this manuscript. This work was supported by the National Science Foundation Doctoral Dissertation Research Improvement Grant (grant number BSC1752393) and the New York University MacCracken Fellowship.

## Region abbreviations (throughout)

A1C: primary auditory cortex;
AMY: amygdala;
CBC: cerebellar cortex;
DFC: dorsolateral prefrontal cortex;
HIP: hippocampus;
IPC: inferior posterior parietal cortex;
ITC: inferior temporal cortex;
M1C: primary motor cortex;
MDT: mediodorsal nucleus of the thalamus;
MFC: medial prefrontal cortex;
OFC: orbital prefrontal cortex;
S1C: primary somatosensory cortex;
STC: superior temporal cortex;
STR: striatum;
V1C: primary visual cortex;
VFC: ventrolateral prefrontal cortex.

## Supplementary Materials for

### Supplementary Figures

**Figure S1.**
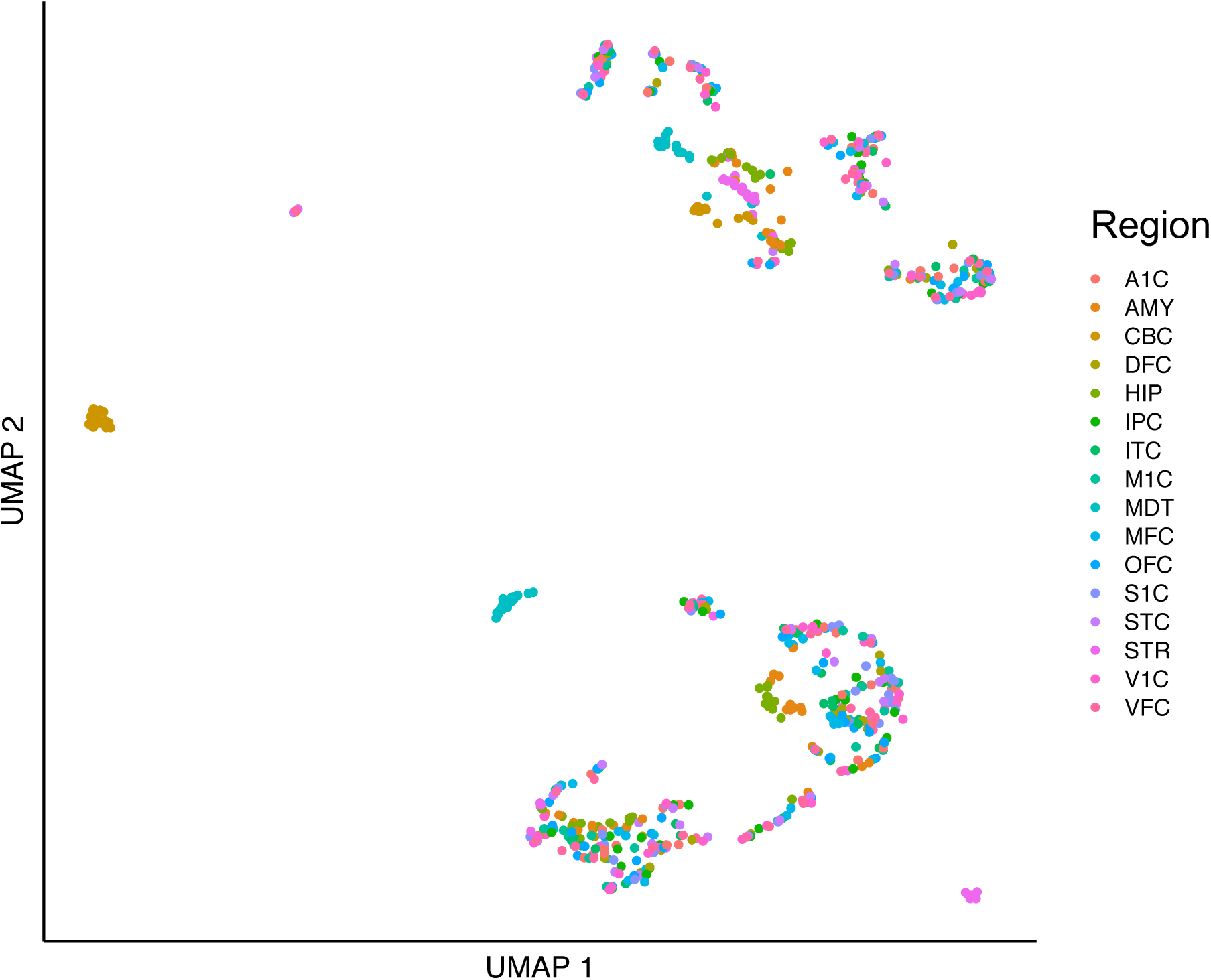
Uniform Manifold Approximation and Projection (UMAP) plot of brain gene expression. Each dot represents one sample. Dot colors indicate brain region (see legend).

**Figure S2.**
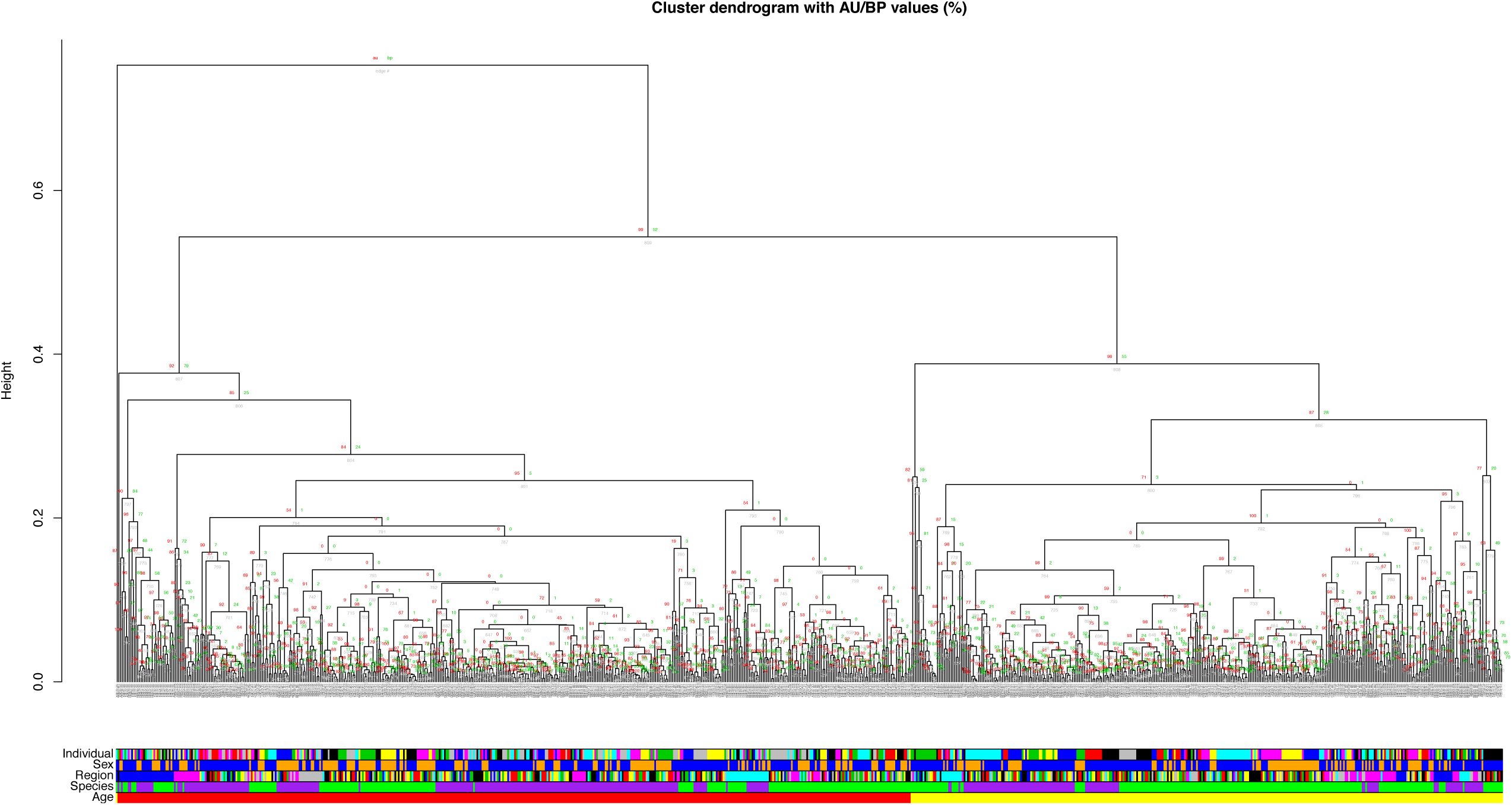
Dendrogram depicting hierarchical clustering across all samples. Each tip represents one sample. AU = approximately unbiased *p*-value; BP = bootstrap probability value. Bottom panels indicate individual, sex (blue = male; yellow = female), region, species (purple = macaque; green = human), and age group (red = fetal; yellow = non-fetal).

**Figure S3.**
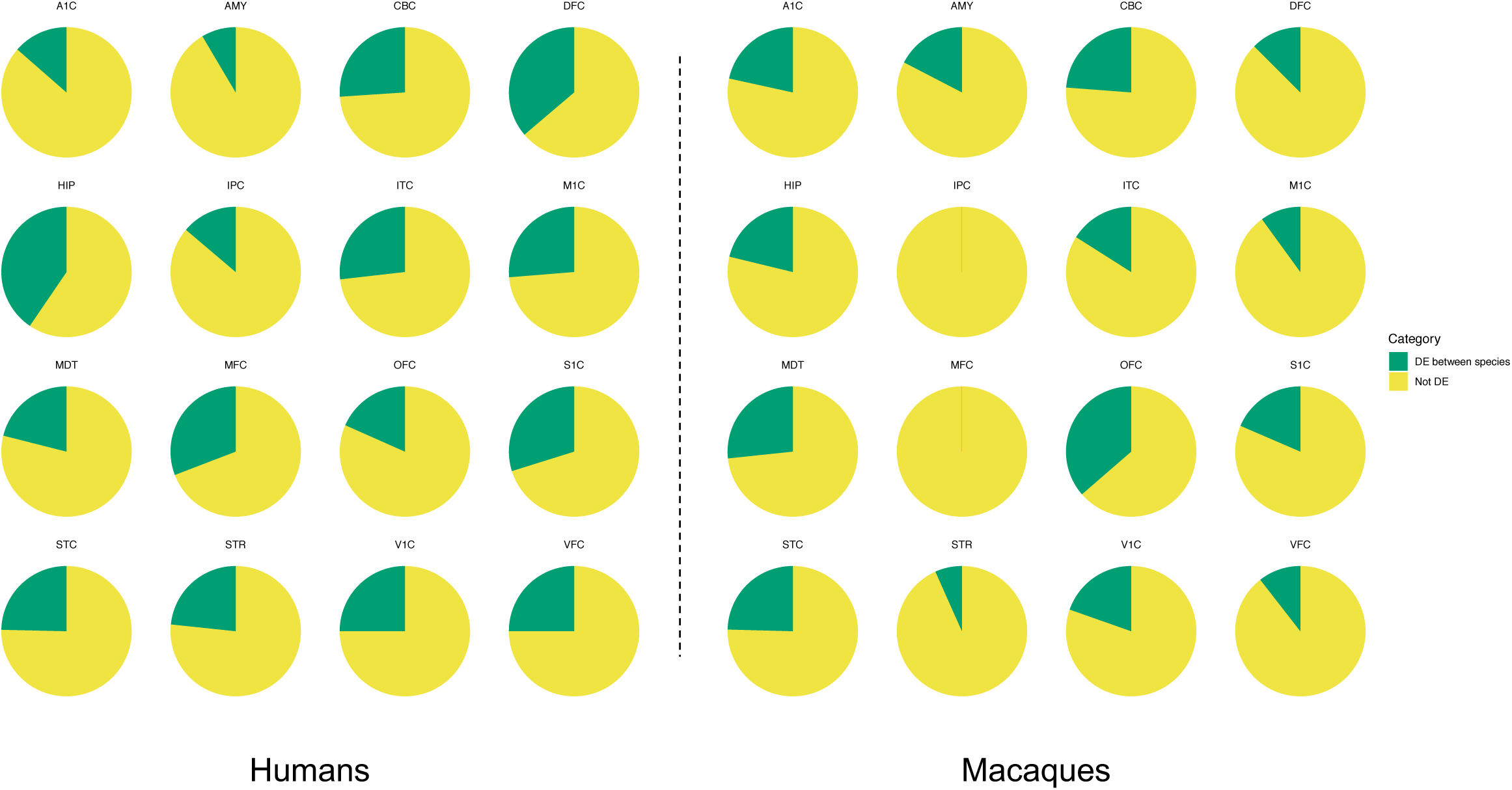
Many sex-biased genes are also differentially expressed between species. Pie charts indicate the proportion of sex-biased genes (identified using the p-value cutoff p<0.005) that are also either differentially expressed (DE) or not DE across species (humans = left panel; macaques = right panel) and regions. Additional results are provided in Tables S2-S4.

**Figure S4.**
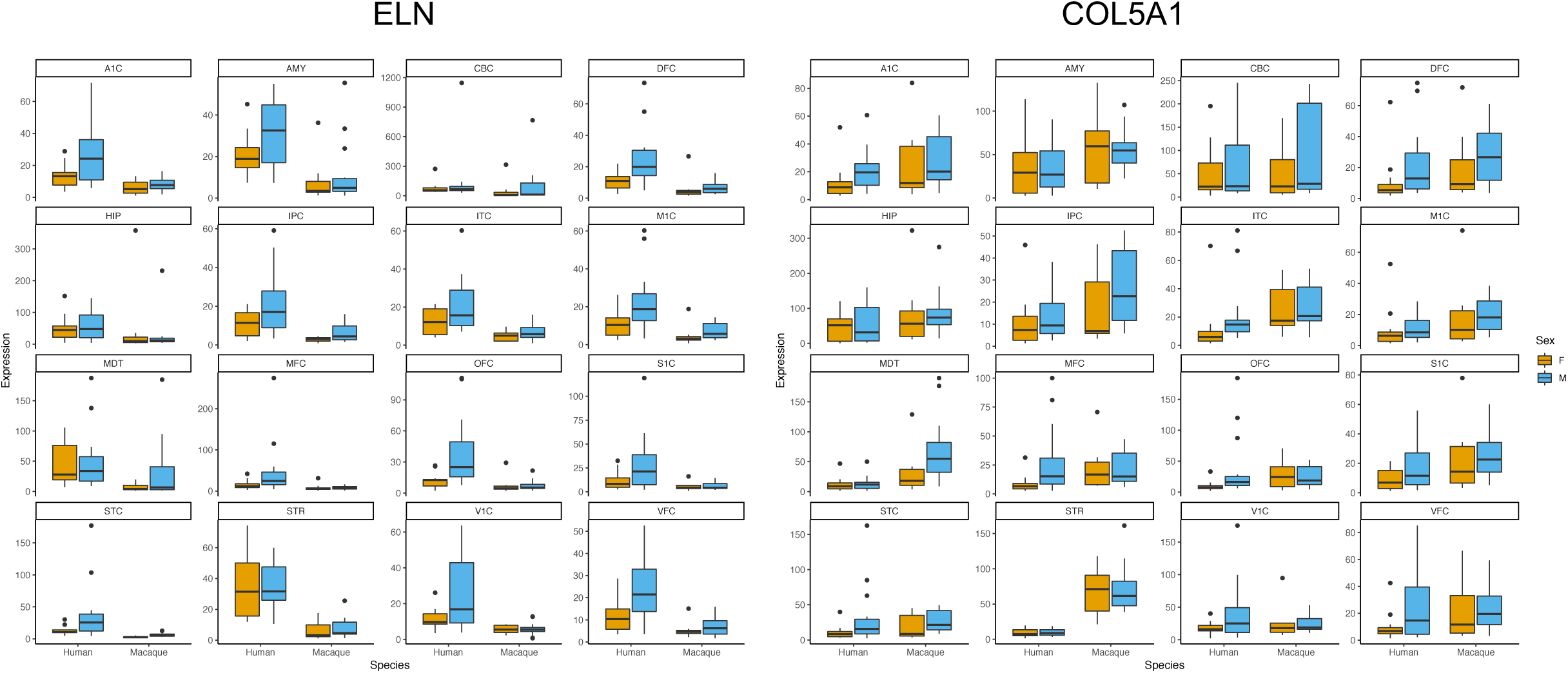
Humans exhibit more extensive male-biased expression of *COL5A1* and *ELN* throughout the brain. Boxplots depict normalized expression levels of *COL5A1* (left) and *ELN* (right) for males and females in each region and species (horizontal line = median, boxes = interquartile range [IQR], whiskers = 1.5*IQR, dots = outlying points). One outlying point (human male, ID=194) was not included here (but was included in the analyses) since high expression values made boxplot visualizations less clear. Colors indicate sex (orange = female; blue = male).

**Figure S5.**
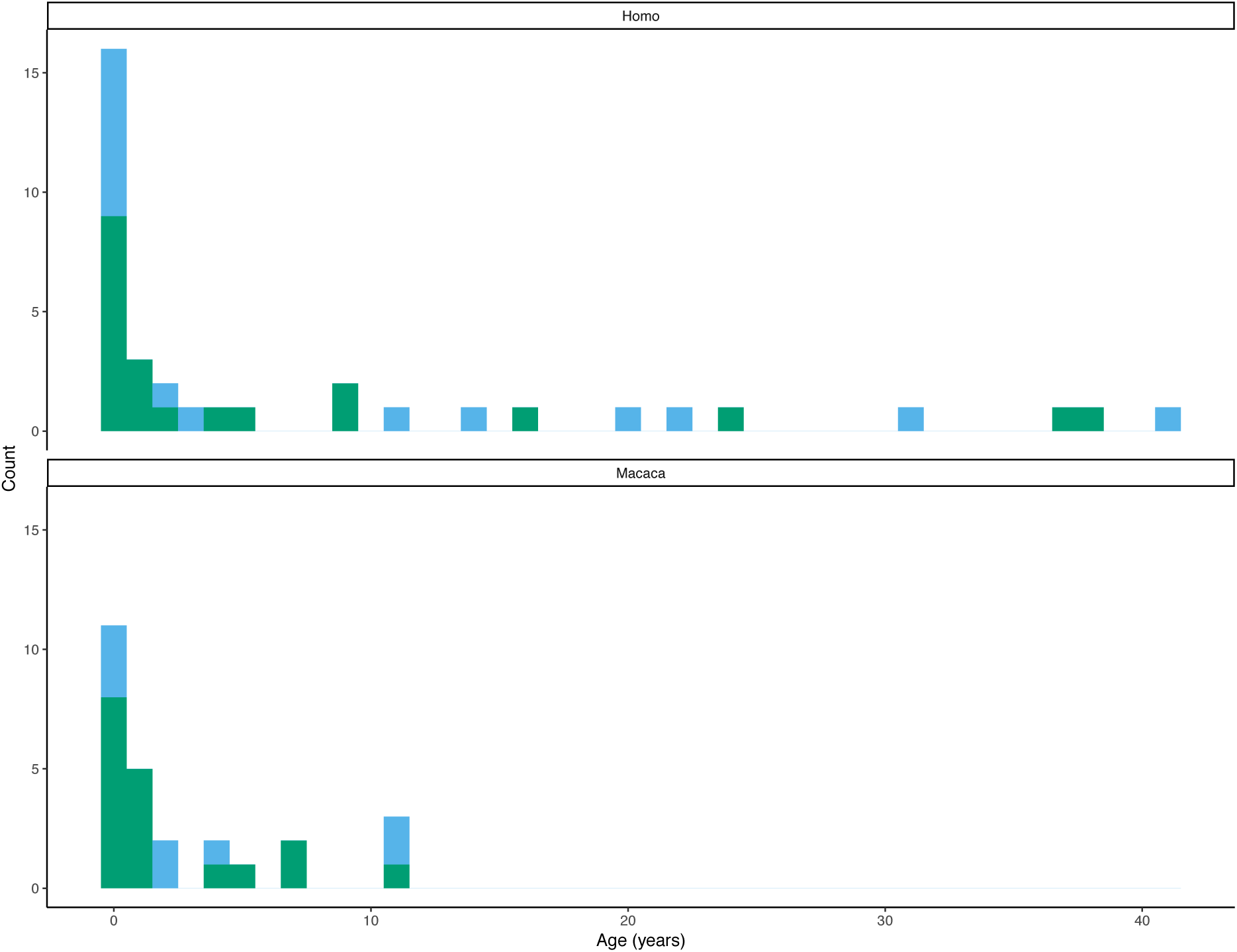
Demographics for individuals included in the source data set. Number of individuals at each age for each species is depicted. Colors indicate sex (blue = female; green = male).

### Supplementary Tables

**Table S1 | Number of differentially expressed genes for each species and region across three p-value cutoffs.**

MB = male-biased; FB = female biased. Results from analyses in which all samples were included are presented in the “All ages” portion of the table. Results from analyses of fetal and non-fetal subsets are presented in their respective portions of the table. Results from sample size permutations (which included both age groups) are represented in the “All ages (median across permutations)” portion of the table. The latter represents the median numbers of sex-biased genes in each category across 100 iterations. Colors indicate significant differences (p<0.05) in the number of male-versus female-biased genes (blue = significantly more male-biased genes; orange = significantly more female-biased genes).

**Table S2 | Percentages of sex-biased genes that are also differentially expressed across species across three p-value cutoffs.**

The “Overall” row indicates the percentages of genes that exhibit sex-bias in at least one brain region and that are also differentially expressed between species in at least one brain region. Rows for specific regions indicate the percentages of genes that both exhibit sex-bias and are differentially expressed between species within that region.

**Table S3 | Percentages of sex-biased genes that are also differentially expressed across species across sample size permutations.**

SB = sex-biased; DE = differentially expressed between species. These are the percentages of genes that exhibit sex-bias in at least one brain region and are also differentially expressed between species in at least one brain region. Results are shown for each of 100 iterations.

**Table S4 | Percentages of sex-biased genes that are also differentially expressed across species within each region across sample size permutations.**

These are the percentages of genes that both exhibit sex-bias and are differentially expressed between species within that region. Results are shown for each of 100 iterations.

**Table S5 | Regression results of sex effects across species for each brain region.**

Coefficient = estimates regression coefficient (negative = genes tend to be biased in opposite directions across species; positive = genes tend to be biased in the same direction across species); P-value = p-value on regression coefficient; R2 = R-squared value for the regression model. Results from analyses in which all samples were included are presented in the “All ages” portion of the table. Results from analyses of fetal and non-fetal subsets are presented in their respective portions of the table. Results from sample size permutations (which included both age groups) are represented in the “All ages (mean across permutations)” portion of the table. The latter represents the mean regression coefficient across 100 iterations.

**Table S6 | Lists of genes exhibiting sex-biased expression (p-value cutoff: p<0.005) in both macaques and humans in each brain region.**

log2FC = log2 fold change between males and females (positive = higher expression in males; negative = higher expression in females). “Direction” indicates direction of bias across species (i.e. male-biased in both species; female-biased in both species; or biased in opposite directions across species). Results from analyses in which all samples were included are presented in the “All ages” portion of the table. Results from analyses of fetal and non-fetal subsets are presented in their respective portions of the table.

**Table S7 | Lists of genes exhibiting sex-biased expression (p-value cutoff: p<0.01) in both macaques and humans in each brain region.**

log2FC = log2 fold change between males and females (positive = higher expression in males; negative = higher expression in females). “Direction” indicates direction of bias across species (i.e. male-biased in both species; female-biased in both species; or biased in opposite directions across species). Results from analyses in which all samples were included are presented in the “All ages” portion of the table. Results from analyses of fetal and non-fetal subsets are presented in their respective portions of the table.

**Table S8 | Lists of genes exhibiting sex-biased expression (p-value cutoff: p<0.05) in both macaques and humans in each brain region.**

log2FC = log2 fold change between males and females (positive = higher expression in males; negative = higher expression in females). “Direction” indicates direction of bias across species (i.e. male-biased in both species; female-biased in both species; or biased in opposite directions across species). Results from analyses in which all samples were included are presented in the “All ages” portion of the table. Results from analyses of fetal and non-fetal subsets are presented in their respective portions of the table.

**Table S9 | Correlations between pairs of regions per species and age group.**

Correlations between sex effects across pairs of regions are listed (negative = genes tend to be biased in opposite directions across regions; positive = genes tend to be biased in the same direction across regions). Results from analyses in which all samples were included are presented in the “All ages” portion of the table. Results from analyses of fetal and non-fetal subsets are presented in their respective portions of the table.

**Table S10 | Differences in tissue specificity between sex-biased and non-biased genes across sample size iterations.**

Results from t-tests comparing mean tissue specificity between sex-biased and non-biased genes are presented for each species across 100 iterations. T-values represent the size of the difference relative to the variation in the data (positive values = non-biased genes exhibit higher tissue specificity; negative values = sex-biased genes exhibit higher tissue specificity).

**Table S11 | Top 30 significant (p<0.05) gene ontology for male- and female-biased genes across species, p-value cutoffs, and age groups.**

Results from human versus macaque analyses are differentiated by the “Species” column (Column B). Results using different p-value cutoffs for sex-biased genes are differentiated in the “Cutoff” column (Column D). Results from analyses in which all samples were included are presented in rows where “All ages” is listed in the “Age Group” column (Column A). Results from analyses of fetal and non-fetal subsets are presented in rows where these group names are listed in the “Age Group” column (Column A). The “Annotated” column lists the number of genes that are annotated with the term. The “Significant” column lists the number of significantly sex-biased genes annotated with that term. The “Expected” column lists the number of genes that would be expected to be sex-biased and annotated with that term under random chance. P-values are from Fisher’s Exact Tests.

**Table S12 | Motif enrichment analyses for sex hormones across species and age groups.**

The “#/% Target w/ Motif” columns indicate the number/percent of sex-biased (aka target genes) with regulatory regions that include the motif. The “#/% Background w/ Motif” columns indicate the number/percent of non-biased (aka background genes) with regulatory regions that include the motif.

**Table S13 | Meta data for samples included in this study (from Zhu et al. 2018).**

Information for each sample, including individual ID, age (in days), age group (fetal vs. non-fetal), sex, species, and batch (i.e. RNA extraction method).

